# The nephronophthisis protein *GLIS2*/*NPHP7* is required for the DNA damage response in kidney tubular epithelial cells

**DOI:** 10.1101/2024.11.02.621644

**Authors:** Lena K. Ebert, Lukas Schloesser, Laura E. Frech, Manaswita Jain, Claudia Dafinger, Max C. Liebau, Thomas Benzing, Bernhard Schermer, Gisela G. Slaats

## Abstract

Nephronophthisis (NPH) is an autosomal-recessive cystic kidney disease representing the most frequent genetic cause of end-stage kidney failure in children and adolescents. NPH is caused by genetic variants in >20 NPHP genes. While nearly all NPHP genes encode ciliary proteins, classifying NPH as a renal ciliopathy, there is evidence for a pathogenic role of a compromised DNA damage response (DDR). Here, we present a novel *Nphp7/Glis2*-deficient mouse model with an early stop codon using CRISPR/Cas9-mediated genome editing (*Glis2*^Y122X^). Homozygous mice develop cystic kidney disease with significant fibrosis at a higher age. Interestingly, the kidneys of these animals exhibit an accumulation of DNA damage (DD) early on, even before any functional impairment of the kidneys becomes apparent. Interactome analysis for GLIS2 revealed an array of DDR-related proteins within the GLIS2 protein complex. Consistent with the *in vivo* data, the knockdown of *Glis2* in kidney epithelial cells led to increased DNA damage. Moreover, supporting the role of GLIS2 in the DDR, we demonstrate that a substantial proportion of GLIS2 is present within the chromatin fraction of cells which is further increased upon UV-induced DD. Live-cell imaging revealed the rapid recruitment of GFP-tagged GLIS2 to sites of laser-induced DD, a response diminished in Glis2^Y122X^ and a variant of Glis2 resembling a known patient mutation. Overall, our data provide compelling evidence for the direct involvement of GLIS2 in the DNA damage response, highlighting the loss of genome stability as an important factor contributing to the pathogenesis of renal ciliopathies.

**Author Summary:** Nephronophthisis (NPH) is a rare inherited kidney disease characterized by cyst formation, fibrosis, and kidney failure at a young age. It is typically caused by mutations in genes essential for the proper function of cilia, small sensory, antenna-like cell protrusions. However, emerging evidence suggests that genome instability and impaired DNA repair may also contribute to NPH. Thus, in some cases, ciliopathies may result from defects in nuclear proteins rather than ciliary proteins. To investigate this, we generated a mouse model with a defective *NPHP7/Glis2* gene using genome-editing tools. The encoded protein GLIS2 has been described to be in the cilium and the nucleus, while its detailed function remained unclear. As these mice aged, they exhibited signs of DNA damage and later developed cystofibrotic kidney disease. Consistently, we demonstrated that the GLIS2 protein interacts with key DNA repair proteins and is rapidly recruited to sites of DNA damage and repair. Thus, inefficient DNA damage repair appears to contribute to kidney disease in this mouse model. These findings underscore the crucial role of genome stability in preventing kidney disease, providing new insights into the underlying causes of NPH.

## Introduction

Nephronophthisis (NPH) is an autosomal-recessive cystic kidney disease representing the most frequent genetic cause of end-stage kidney failure in children and adolescents [1]. NPH can be caused by mutations in NPHP genes, and since the identification of NPHP1 in 1997 [2], over 20 additional NPHP genes have been identified [1,3–5], however, not all clinical cases can be explained by variants in those genes [6,7]. Almost all NPH-associated genes encode for proteins localizing to primary cilia. In the kidney, these sensory organelles project from the apical site of kidney tubular epithelial cells into the lumen of the tubules. Cilia are present in the vast majority of mammalian cell types which can explain the broad spectrum of extrarenal phenotypes in NPH and related renal ciliopathies [8]. The importance of cilia in the pathogenesis of cystic kidney diseases has long been established. However, there is also evidence for extraciliary functions of some ciliopathy proteins. Here, their involvement in the DNA damage response (DDR) signaling is particularly interesting: With *ZNF423*/*NPHP14* and *CEP164*/*NPHP15*, disease-causing mutations in genes implicated in DDR have been identified in NPH patients [9]. In addition, SDCCAG8/NPHP10 [10], CEP290/NPHP6 [11], MAPKBP1/NPHP20 (12) and NEK8/NPHP9 [12] have been linked to DNA replication stress and DNA damage signaling, thereby suggesting a role for DDR signaling in the pathogenesis of renal ciliopathies. However, a direct role of proteins encoded by NPHP genes in the DDR remained to be investigated [13].

Here, we focus on *GLIS2/NPHP7*, a gene encoding for GLIS family zinc finger protein 2, which acts as a transcription factor [14]. GLIS2 is essential for maintaining kidney function and regulates the expression of genes involved in inflammation, fibrosis, senescence, and cell death [15–17]. The protein is characterized by five zinc finger domains [18]. To date, only three human mutations causing NPH-related ciliopathies have been identified: The first *GLIS2* mutation described is a transversion (c.775+1G>T, ClinVar ID: 242364), initially reported as IVS5+1G>T, which abrogates the 5 prime obligatory donor splice site of coding exon 5 [15]. Another homozygous *GLIS2* missense mutation (c.523T>C, p.C175R, ClinVar ID: 127160) was identified in a large NPH-related cohort [19]. More recently, a novel homozygous in-frame deletion in *GLIS2* (c.570_584del, p.N190_V194del, ClinVar ID: 1299703) was identified using whole exome sequencings (WES) [20]. Additional evidence for the impact of GLIS2 in kidney maintenance comes from two classical knockout mouse models that resemble some of the histopathological features of NPH [15,16]. Here, we present a novel *Glis2*-deficient mouse model (*Glis2*^Y122X^) in which we inserted a stop codon with CRISPR/Cas9-mediated genome editing. Homozygous mice display a late-onset and slowly progressing cystic-fibrotic kidney disease with a pronounced accumulation of DNA damage, which precedes cyst formation and the loss of kidney function. We further connect GLIS2 to the DDR by protein interactome and chromatin binding analyses and demonstrate the recruitment of GLIS2 to the place of DD inside the nucleus.

## Results

### A novel *Glis2*^Y122X^ mouse model resembles NPH

Using CRISPR/Cas9-mediated genome editing, we generated a novel *Glis2* mutant mouse line (Fig. 1A): An asymmetric single-stranded donor oligonucleotide (ssODN) was used to introduce a single point mutation in exon 4 (coding exon 3), resulting in an early stop codon, referred to as *Glis2*^Y122X^ (c.366T>A, p.Y122X) which was confirmed by sequencing both genomic DNA and reverse-transcribed cDNA (Suppl. Fig. 1A/B). The stop codon is located upstream of the five zinc finger domains of GLIS2, which are encoded by exons 4-7. Homozygous *Glis2*^Y122X^ animals were born in lower numbers than expected by Mendelian ratio (Suppl. Fig. 1C), while Glis2 mRNA abundance was not significantly altered (Suppl. Fig. 1D). Histological analysis of kidney tissue from homozygous *Glis2*^Y122X^ mice at various ages revealed subtle alterations, including some dilated tubules and mild tubular basement membrane disintegration with irregular thickening by 16 weeks of age (Fig. 1B). At one year of age, distinct small cysts as well as immune infiltrates were evident, and these features were significantly more pronounced with increasing age with severe cyst formation in *Glis2*^Y122X^ mice of > 1.5 years. In comparison to previously published *Glis2* knockout mice [15,16], this new *Glis2*^Y122X^ model displayed slower disease progression and less pronounced medulla region degeneration. As for kidney function, serum urea values of *Glis2*^Y122X^ mice showed a slight increase at the age of 16 weeks and were significantly elevated at higher age as compared to littermate control animals (Fig. 1C/D). Consistent with renal fibrosis, mRNA expression analysis done by qPCR of RNA from total kidneys for a panel of senescence, fibrosis and inflammatory response markers revealed transcriptional upregulation of *Col1a1*, *Ctgf*, *Tgfb1*, *Tnfa*, and *Il1b* in *Glis2*^Y122X^ mice at the age of 16 weeks and >12 months (Fig. 1E/F). Taken together, the clinical and histological phenotype of *Glis2*^Y122X^ mice with cyst formation, interstitial fibrosis and chronic kidney disease mirrors several key features of human NPH.

**Fig. 1:**
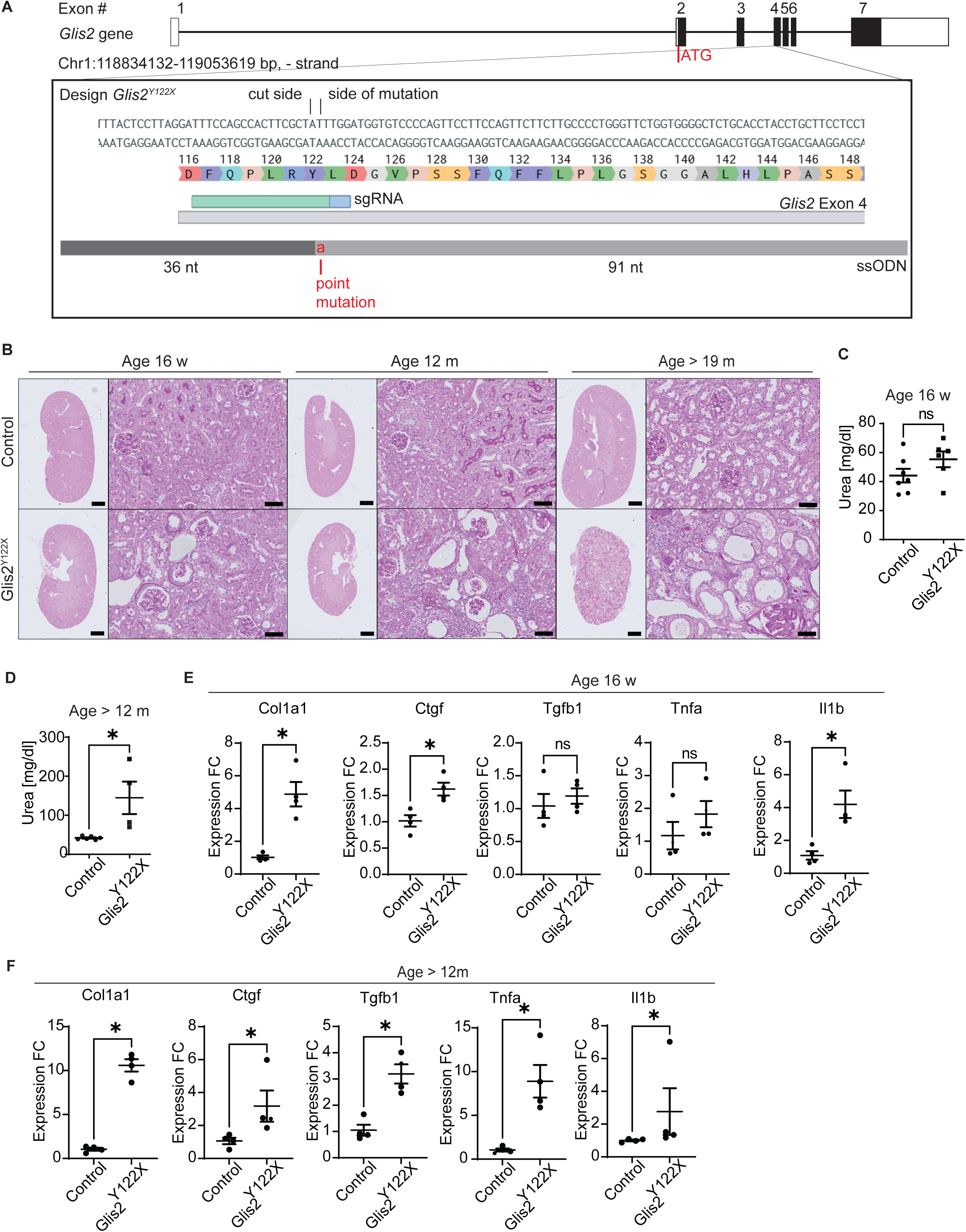
The novel mouse model *Glis2*^Y122X^ demonstrates kidney characteristics of NPH. (a) Strategy to generate the *Glis2*^Y122X^ mouse model. On the top the exon (boxes)-intron (lines) structure of *Glis2* is indicated. This model was generated using one sgRNAs targeting the beginning of exon 4 (coding exon 3) combined with a ssODN introducing a point mutation which leads to a stop codon. (b) Periodic acid-Schiff (PAS) staining of *Glis2*^Y122X^ and control mice at 16 weeks, 12 months and >19 months of age. (scale bars: overview pictures 1000 µm, detailed pictures 50 µm, N=5 (16w), 3 (12m), 2 (>19m). (c) Serum urea levels of 16 week old *Glis2*^Y122X^ and control mice (N=6). (d) Serum urea levels of >12 months old *Glis2*^Y122X^ and control mice (N >=4). (e,f) Quantitative real-time PCR of indicated marker genes for fibrosis, senescence, and inflammation showing upregulation in kidney tissue of 16 week old (e) and >12 months old (12-20 months old) (f) *Glis2*^Y122X^ mice compared to age matched control mice. Indicated genes were normalized to *Hrpt*. Statistical analysis was performed by a unpaired Student’s 2-tailed t-test with a p-value < 0.05 considered significant (N=4).

### The loss of wildtype *Glis2* expression leads to increased DNA damage

Consistent with studies that suggested a role for NPHP proteins in DNA damage response signaling [9–13], the kidneys of *Glis2*^Y122X^ mice at an age of 16 weeks exhibited increased TUNEL positivity (Fig. 2A) and an elevated signal for phosphorylated H2AX (γH2AX) an established marker for DNA strand breaks (DSB; Fig. 2B). This increase occurred at a stage when only mild histological alterations were observed, while kidney function remained largely unaffected, supporting the notion that it was not a consequence of kidney failure or elevated uremic toxins but rather the result of an intrinsic defect. Consistently, γH2AX elevation was not detected in extrarenal tissues, as demonstrated for the liver (Fig. S1E). To confirm the accumulation of DD and DSBs in relation to *Glis2* by independent means, we targeted *Glis2* in murine kidney epithelial cells (murine inner medulla collecting duct cells (mIMCD3)) using siRNA (Fig. 2C) and determined the levels of DNA fragmentation by neutral comet assays. Upon knockdown of *Glis2*, cells showed significantly more DSBs compared to controls with a more pronounced effect in the presence of the DNA replication inhibitor aphidicolin (Fig. 2D)[21]. Taken together, these results show increased DSB and accumulation of DD in the *Glis2*^Y122X^ mice and upon reduction of *Glis2* in mIMCD3 cells.

**Fig. 2:**
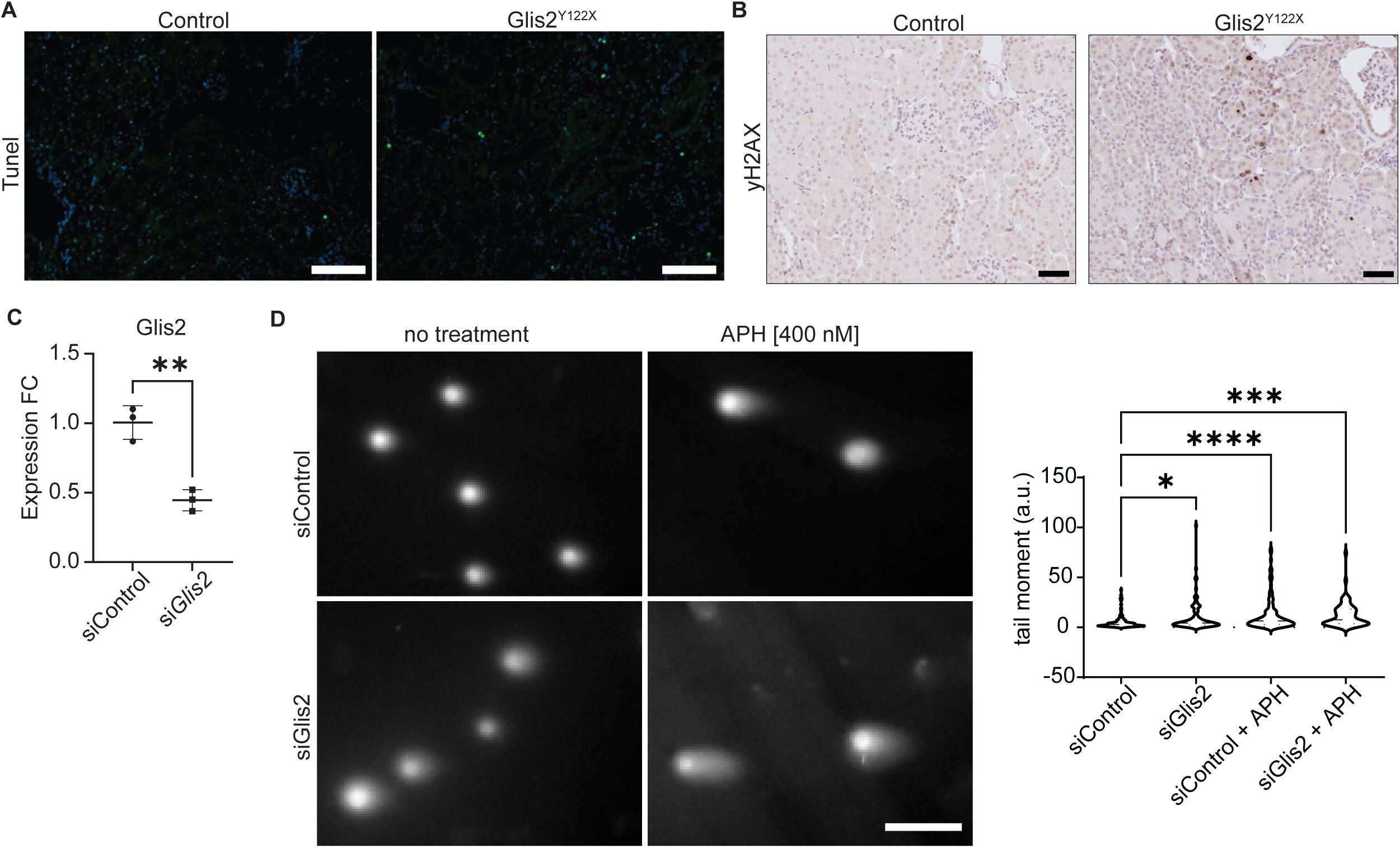
*Glis2*^Y122X^ mice display accumulation of DNA damage. (a) Tunel staining and (b) yH2A.X staining of renal tissue of 16-week-old *Glis2*^Y122X^ and control mice (scale bars: tunel 100 µm, yH2A.X 50 µm). (c) Quantitative real-time PCR of *Glis2* to validate the siRNA knockdown (N=3). (d) Representative images of *Glis2*-depleted and control mIMCD3 cell DNA stained with SYBR Gold nucleic acid gel stain after performing a neutral comet assay with or without aphidicolin (APH) treatment (400 nM, 18 hours, scale bar = 100 μm). At least 30 comets were scored per condition per experiment to evaluate the comet tail moment by using one-way ANOVA, Tukeýs multiple comparisons test (N = 3, \**P* < 0.05, \*\*\**P* < 0.001).

### The GLIS2 protein complex contains DDR key players

To gain molecular insights into the mechanisms underlying the role of GLIS2 in DDR signaling we analyzed whether GLIS2 is associated with known DDR proteins in a common protein complex. To address this question, we transiently expressed either FLAG-tagged GLIS2 (FLAG.GLIS2) or GFP (FLAG.GFP) in HEK293T cells. Pull-down assays with anti-FLAG antibodies were performed and analyzed by mass spectrometry to identify specific components of the GLIS2 protein complex. Principle component analysis revealed that the samples clustered together within their specific groups (FLAG.GFP and FLAG.GLIS2) and that the groups were separated by the largest variance (Fig. 3A). A two-sample t-test identified many proteins (926) significantly enriched upon pulldown of FLAG.GLIS2 as compared to the FLAG.GFP control (Fig. 3B). As expected, GLIS2 was among the most enriched proteins, confirming the successful immunoprecipitation of FLAG.GLIS2. Gene Ontology (GO) term enrichment analysis of the interactome showed cellular compartments “nuclear speckles”, “nucleolus” and “nucleoplasm” to be the most enriched (Fig. 3C). Among the significantly enriched proteins, we identified several proteins known to be involved in DDR and to be dynamically recruited to the sites of DNA damage. These included Bcl-2-associated transcription factor 1, a radiation-induced interactor of H2AX [22], the DNA damage detector protein Poly[ADP-ribose]polymerase 1 (PARP1) [23,24] as well as several factors modulated or recruited by PARP1 to DNA breaks such as the Chromodomain-helicase-DNA-binding protein 1-like (CHD1L) [25] and Chromodomain-helicase-DNA-binding protein 4 (CHD4) that are involved in chromatin remodeling at DNA breaks [26], ATP-dependent RNA helicase DDX3X [27], the Double-strand break repair protein MRE11A [28] or the Proliferating cell nuclear antigen PCNA [29](Fig. 3B/D, Suppl. Table 2). The presence of these DDR proteins within the complex of GLIS2 together with the accumulation of DSBs in GLIS2 deficient tissue and cells supported the idea of Glis2 being directly involved in DNA repair pathways.

**Fig. 3:**
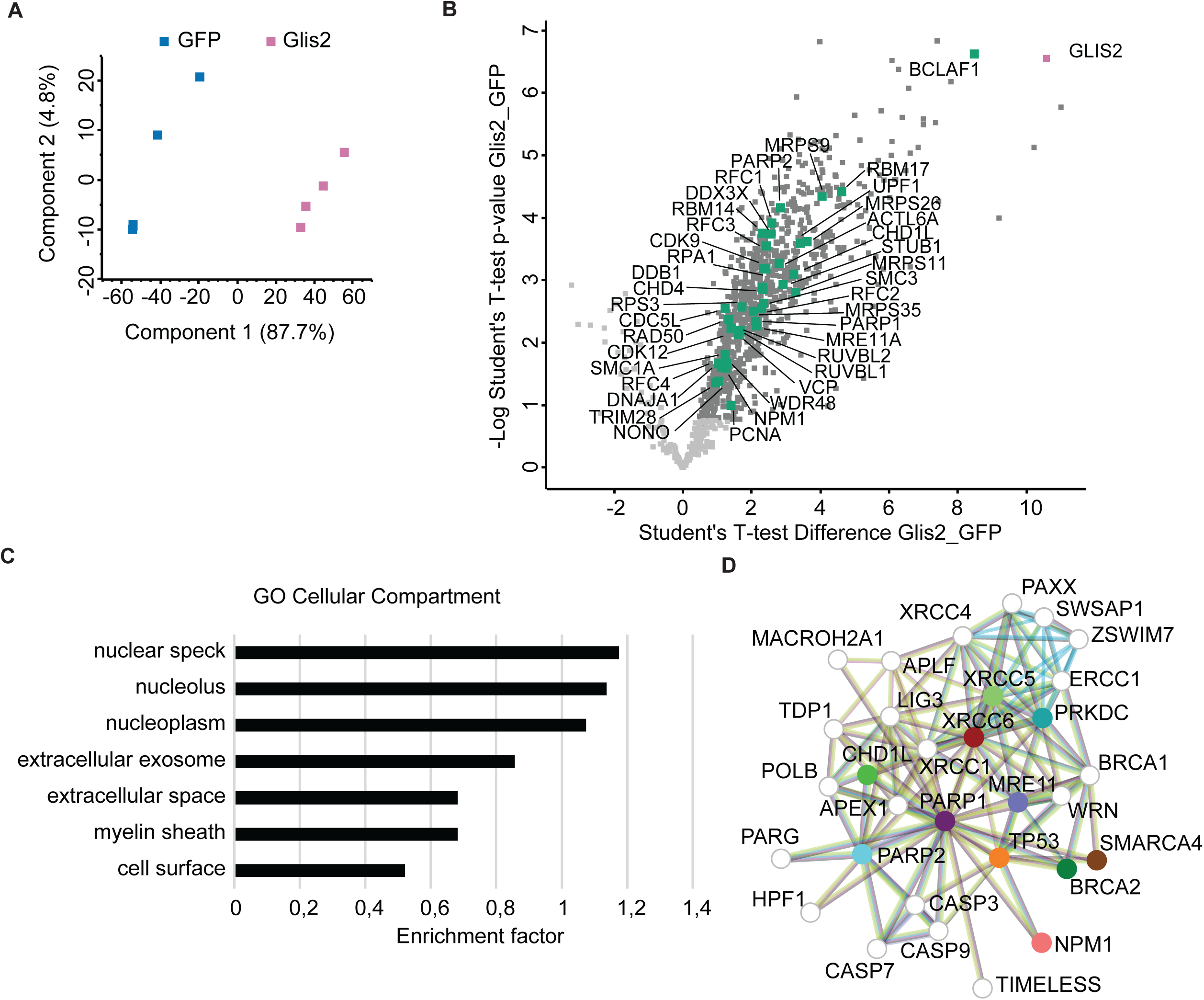
The GLIS2 interactome reveals a connection to DNA damage response. (a) Principal component analysis scatter plot of FLAG.GLIS2 and FLAG.GFP. Depicted are the first two principal components (PC1 and PC2). The axes represent the percentages of variation explained by the principal components. (b) Scatter plot with the t-test differences of label-free quantification (LFQ) intensities of the FLAG.GLIS2 pulldown vs FLAG.GFP pulldown on the x-axis and the statistical significance (-log_10_ Student’s t-test p-value) on the y-axis. Highlighted in dark grey are proteins that are significantly regulated based on the Student’s t-test (s0 = 0, FDR = 0.05). GLIS2 is among the most enriched proteins (pink). Proteins associated with the gene ontology biological process terms „DNA Damage Response, detection of DNA damage” and „DNA repair”, as well proteins discussed in the manuscript are highlighted in green. (c) Gene Ontology term enrichment analysis for the category Cellular Compartments. Significantly enriched terms were selected based on a Fisher’s Exact Test with a FDR 0.02. (d) STRING interaction network of PARP1. Colored notes correspond to significant interactors of GLIS2.

### GLIS2 is recruited to sites of DNA damage

Since the interactome analyses revealed GLIS2 to be associated with numerous proteins involved in DDR, we hypothesized that GLIS2 might be dynamically bound to chromatin and recruited to sites of DNA damage. We therefore performed chromatin fractionation in HEK293T cells which revealed that GFP.GLIS2 was bound to chromatin. Notably, this association was enhanced upon DD induction by UV-C treatment (Fig. 4A). To investigate whether GLIS2 itself is recruited to sites of DD [30], we performed laser irradiation experiments. Live cell imaging of HEK293T cells transiently overexpressing GFP-tagged GLIS2 proteins revealed recruitment of GLIS2 at the sites of DD induced by 266 nm (Fig. 4B) or 800 nm lasers (Fig. 4C/D and Suppl. Movie 1-2). To understand whether alterations in this recruitment might be relevant to disease development, we conducted the same experiments using truncated GLIS2 variants. These variants corresponded to our mouse model or to a mutation observed in patients [15]. For this we used TALEN-based genome editing to create stable polyclonal HEK293T cell lines expressing either GFP-tagged wildtype GLIS2 or two GLIS2 truncations. In the first mutant cell line we introduced the published disease-causing point mutation c.775+1G>T (GFP.GLIS2[IVS5+1G>T]) affecting an obligatory donor splice site [15]. In the other cell line, we expressed a truncated version of the protein expressing only the first 121 N-terminal amino acids (GFP.GLIS2[1–121]), thus lacking all zinc finger motives (Suppl. Fig. 2A-C). We tested the capacity of the different GLIS2 proteins to recruit to sites of 800 nm laser-induced DNA damage. The stable GFP.GLIS2 cell line confirmed our results using transiently expressed GLIS2 (Fig. 4B-D) and was always recruited to the site of damage (Fig. 4E). Both mutant cell lines showed a reduced recruitment to the place of DD after laser ablation. While GFP.GLIS2[IVS5+1G>T] showed reduced recruitment to the DD site, GFP.GLIS2[1–121] lost this recruitment almost completely (Fig. 4E, Suppl. Fig. 2C).

**Fig. 4:**
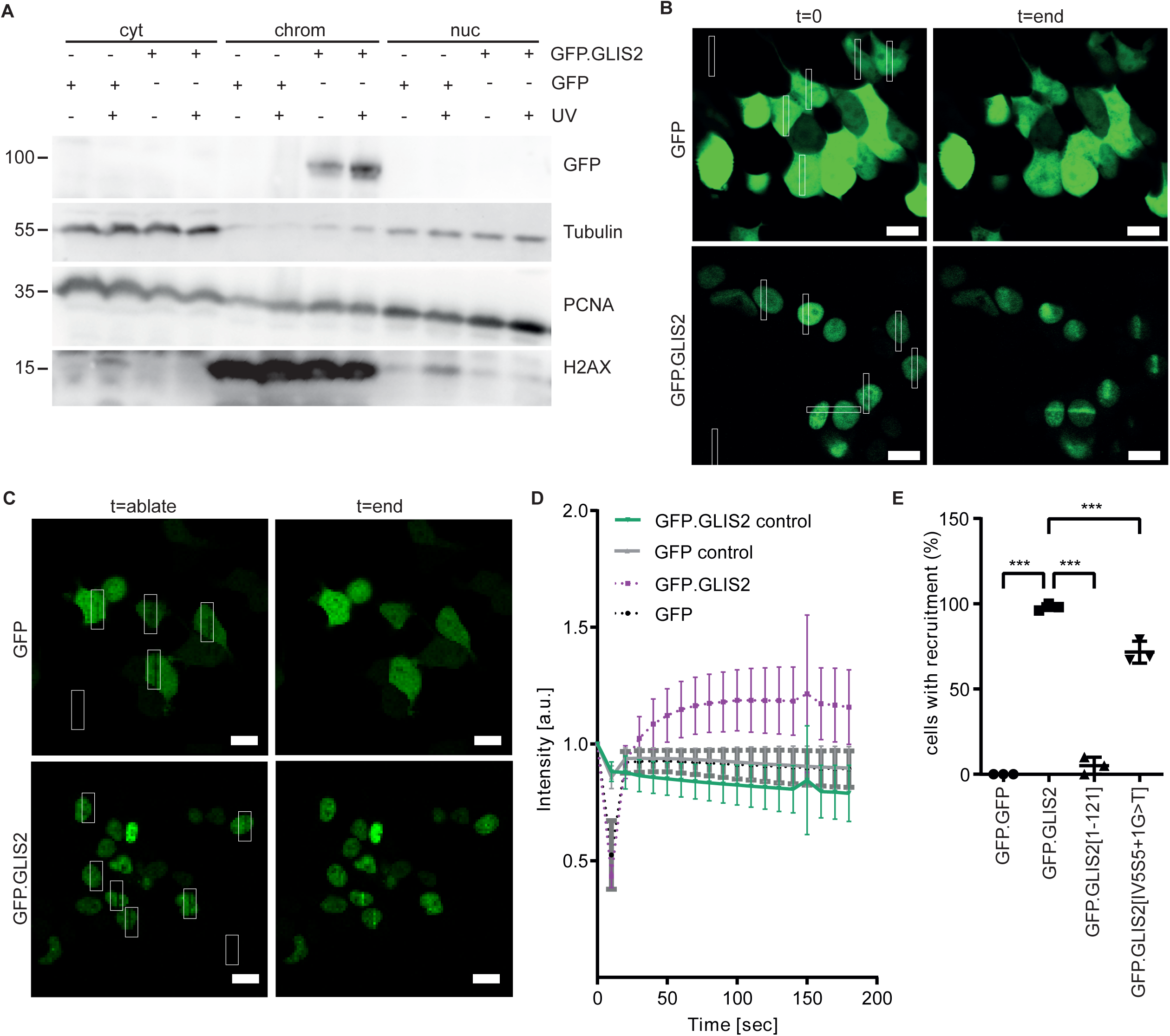
GLIS2 is recruited to the sites of DNA damage. (a) Western blot of chromatin fractionation of GFP and GFP.GLIS2 overexpressing HEK293T cells. GFP.GLIS2 is not expressed in the cytosol (tubulin) or nuclear (PCNA) fraction, but in the chromatin-enriched fraction (H2AX). Samples were run on parallel gels contemporaneously (N=3) (b) GFP-tagged GLIS2 transfected HEK293T cells were ablated by 266 nm laser irradiation (white squares, t = 0 s). Redistribution of GFP.GLIS2 was followed over time (end t >45 s), scale bar = 10 µm (N=3). (c) GFP–tagged GLIS2 transfected HEK293T cells were ablated by 800 nm laser irradiation (white squares, t = 10 s (ablation)). Redistribution of GFP.GLIS2 was followed over time (end t = max 180 s, scale bar = 10 µm; N=3). (d) Quantification of the GFP (black) or GFP.GLIS2 (violet) recruitment upon 800 nm laser irradiation. Regions outside of cells were measured as controls (grey and green). The intensity at t=0 was normalized to 1. Approx. 25 cells were analyzed per replicate (N=3). (e) Quantification of the percentage of ablated cells that showed GFP.GFP or GFP.GLIS2 wildtype or variant recruitment upon 800 nm irradiation. At least 52 cells have been analyzed per replicate and cell line (N=3).

## Discussion

This study was motivated by the fact that there are currently only a few mouse models that capture key aspects of the human ciliopathy NPH, and that CRISPR/Cas9-based genome editing has significantly improved the efficiency of generating knockout mice. Therefore, we aimed to generate a model for nephronophthisis by introducing a targeted point mutation in exon 4 (3^rd^ coding exon). Surprisingly, the homozygous knockout animals exhibit a much later phenotype compared to the previously known classical knockout models: One of these previous models, with a deletion from exon 3 to exon 5, shows severe impairment of corticomedullary differentiation at 4 weeks and, subsequently, a reduction in the medulla region and kidney size [15]. Another model, created around the same time with a deletion of exon 6, shows initial changes at 4 months and by 7 months presents with shrunken and atrophic kidneys and significantly reduced survival [16]. In contrast to these knockout mice, our model with the stop codon in exon 4, *Glis2*^Y122X^, at 16 weeks of age shows a very mild phenotype with dilated tubules without significant changes in kidney function. This intensifies at 12 months and older, with massive fibrosis and kidney failure. Despite the obvious differences in the timeline of kidney pathology, all three models suggest that the loss of Glis2 does not have a major impact on kidney development, but rather affects the adolescent/adult kidney. This is consistent human nephronophthisis caused by *NPHP7* mutations [15,19].

In our *Glis2*^Y122X^ model, a phenotype only appears in adulthood, suggesting that the absence of GLIS2 causes cumulative damage in the adult kidney, followed by fibrosis. GLIS2 is a known transcription factor that directly binds DNA with its zinc finger domains [14]. Furthermore, an initial study suggested a ciliary localization for GLIS2 [15], which later studies and our experiments with GFP-tagged GLIS2 have not yet been able to confirm. Notably, we found evidence of DNA damage accumulation in the kidneys of homozygous animals at an early stage, before the onset of fibrosis and kidney failure. To further investigate the molecular functions of GLIS2, we analyzed the interactome of GLIS2 using mass spectrometry. Strikingly, this revealed an association of GLIS2 with numerous key components of DDR signaling, particularly of factors recruited to the site of DNA damage. Notably, PARP1 stands out, which in turn recruits other factors like CHD1L, CHD4, DDX3X, MRE11a or PCNA to the damage site [23,25–29,31]. All these proteins co-precipitated with GLIS2, marking them as components of the GLIS2 protein complex. This led us to hypothesize that the nuclear protein GLIS2, in addition to its role as a transcription factor, might also be involved in DNA damage response signaling. Our live-imaging experiments confirmed this hypothesis: double-strand breaks (DSBs) were induced with a laser, and GLIS2 was rapidly recruited to the site of damage in the nucleus, demonstrating a completely new function for this important protein. It might be that in the context of DNA damage GLIS2 acts as transcriptional repressor [32,33]. Shutting down transcription is an early event in the DDR, facilitated by DDR master regulators like PARP1. Whether GLIS2 is transiently recruited to condense the chromatin structure and cause transcriptional silencing around lesion sites remains to be confirmed. It seems unlikely that GLIS2 functions as an essential DNA damage repair factor. If GLIS2 had a crucial DNA damage repair function, we would expect either progeroid or cancer phenotypes in the mouse models. It is important to note that the loss of GLIS2 primarily manifests in the kidney, with only limited evidence suggesting extrarenal functions. While we did not observe cancer in *GLIS2*^Y122X^ even in high age, renal fibrosis could be regarded as a premature aging phenotype. Besides transcriptional repression, GLIS2 could have a function in DNA damage detection or signaling after recognition of damage, which it would share with several candidates from the interactome.

The involvement of GLIS2 in DDR and the subsequent loss of genomic stability upon GLIS2 loss explain both the late but severe onset of kidney fibrosis and add an important aspect to the recent discussion on whether GLIS2 could be a therapeutic target in autosomal dominant polycystic kidney disease (ADPKD). It has recently been demonstrated that both the loss and the antisense-mediated inhibition of *GLIS2* negatively impact cyst formation in PKD models [34]. This complements an earlier study showing a similarly positive effect in a cystic kidney model resulting from the deletion of *Kif3a* and, thereby, functional cilia [17]. Whether GLIS2 can actually be used as a therapeutic target in ADPKD will depend on how severely the genomic instability associated with GLIS2 inhibition affects the homeostasis of the kidney and extrarenal tissues.

There is approximately less than 5% of treatment available for the more than 7,000 orphan diseases so far identified, most of which have genetic causes [35]. Despite the identification of several candidate drugs in rodent NPH models, there has been a lack of clinical trials, and there is currently no specific therapy that halts disease progression in NPH patients [36]. The work presented here suggests that DDR might be a potential therapeutic target for NPH with the aim of boosting the efficiency of repair and decreasing accumulating damage. Interestingly, PGE1 treatment of *Nphp1*^-/-^ mice has recently been found to improve kidney phenotypes. Transcriptomic analyses revealed alterations in several pathways to be blunted by the treatment, including the DNA damage response [37]. In summary, we hypothesize that any intervention that increases the efficiency of DDR and resolves DD might have the potential to slow the progression of kidney degeneration. This includes dietary interventions, pharmacological treatments with small molecules or possibly with senolytics, epigenetic approaches, or even hormetic stressors. Therefore, interventional studies in suitable mouse models, including the model from this study, are needed to identify and further evaluate appropriate strategies.

## Materials and Methods

### Ethics statement

All matings and experimental protocols in this study were performed in accordance with European, national, and institutional guidelines, and approved by LANUV NRW (Landesamt für Natur, Umwelt und Verbraucherschutz Nordrhein-Westfalen/ State Agency for Nature, Environment and Consumer Protection North Rhine Westphalia.

### Animal models

Mice were housed according to standardized specific pathogen-free conditions at the *in vivo* Research Facility of the CECAD Research Center, University of Cologne, Germany. Mice were kept in individually ventilated cages under a 12-hour light/dark cycle and had free access to food and water. Experiments were conducted on mice with a pure C57BL/6N background. Mice of both sexes were included in this study. The age of the mice used for the experiments is indicated in the figures. Ear biopsies were used for DNA isolation as previously described [38]. For genotyping PCRs REDTaq® ReadyMix™ (Sigma-Aldrich) was used according to the manufacturer’s instructions and visualized via gel-electrophoresis. Primer sequences are listed in Suppl. Table 1.

### Generation of *Glis2*^Y122X^ mice

The *Glis2*^Y122X^ mice where generated in the Transgenic Core Unit of the CECAD Research Center, University of Cologne, Germany). The sgRNAs (Suppl. Table 1) were generated by *in vitro* transcription with a T7 RNA polymerase (NEB) from the pSpCas9(BB)-2A-Puro (PX459) V2.0 plasmid gifted from Feng Zhang (Addgene plasmid # 62988; http://n2t.net/addgene:62988; RRID:Addgene_62988) [39]. sgRNAs were column purified (Qiagen). The custom single-stranded donor oligonucleotide (ssODN, Suppl. Table 1) was ordered from IDT and resuspended in nuclease-free H20 to a concentration of 10 µM. sgRNA and ssODN were stored at −80°C. Mouse transgenesis was performed as previously described using a pronuclear injection (PNI)-based approach [40,41]. In brief, zygotes were collected from the oviduct of superovulated females and inspected for the presence of two pronuclei. Microinjection of the PNI reaction mix containing 50 ng/μL sgRNA, 50 ng/µl Cas9 mRNA (TriLink Biotechnologies), 30 µg/µl Cas9 protein (PNA Bio Inc), and 100 ng/µl ssODNs (IDT) was performed with an Axio Observer.D1 microscope (Zeiss) and microinjector devices CellTram and FemtoJet with TransferMan NK2 micromanipulators (Eppendorf) into the male pronucleus using with injection capillaries (BioMedical Instruments, BM100F-10; type PI-1.6). After overnight cultivation, embryos that developed to the two-cell-stage were transferred into oviducts of pseudo-pregnant foster females. For sgRNA design, sequencing analysis and alignments (including Fig. 1A) the Benchling software (https://benchling.com) was used.

### Serum measurements

Blood samples collected from all mice were kept at room temperature for 30 min and then centrifuged at 1000 rcf for 10 minutes to obtain serum samples. Serum urea levels were measured by the Institute of Clinical Chemistry, University Hospital of Cologne, Germany.

### Renal histology

The kidneys of the mice were harvested at indicated time points and fixed in 4% formalin overnight at 4°C. The tissues were further processed for paraffin embedding at the CECAD Research Center, University Hospital, Cologne, Germany. One µm sections were cut from the paraffin embedded tissue blocks. For morphological analysis, periodic acid-Schiff (PAS) staining was performed as described previously [42]. Tunel Staining was performed with the DeadEnd™ Fluorometric TUNEL System (Promega) following the manufacturer’s instructions, with the exception of incubation with Hoechst (ThermoFisher, 1:1000) for the nuclear staining prior to mounting with ProLong™ Diamond (ThermoFisher Scientific). Images were acquired using the AxioObserver microscope with an axioCam ICc 1, Axiocam 702 mono, Apotome system (Carl Zeiss MicroImaging, Jena, Germany; objective Plan-Apochromat 20x/0.8). Phosphorylated H2A.X staining was performed as described previously using a phospho-Histone H2A.X (Ser139) antibody (Cell Signaling, #2577) [43].

### qPCR

Total RNA was extracted from kidney tissue using TRIzol (Sigma/Invitrogen). One µg of total RNA was used as a template to generate cDNA (ABI High Capacity cDNA Reverse Transcription kit). qPCR was performed with SYBR Green qPCR mix (Applied Biosystem). All qPCR experiments were performed on either the QuantStudio 5 or the 7900HT (both Applied Biosystems). The comparative Ct method was used to perform qPCR analysis. Either *Hrpt* (*in vivo* data) or *β-Actin* (siRNA validation) were used as housekeeping genes, after showing uniform expression with low variance across the sample set. All primer sequences are provided in Suppl. Table 1.

### Cell culture

Immortalized mouse inner medullary collecting duct (mIMCD3) cells were cultured in DMEM:F12 (Sigma) supplemented with GlutaMAX (Gibco) and 10% FBS at 37°C, 5% CO2. HEK293T cells were cultured in DMEM (GlutaMAX, Gibco) supplemented with 10% FBS at 37°C, 5% CO2. Mycoplasma contamination was excluded by regularly performing PCR-based tests (Promokine detection kit).

### Molecular cloning

AAVS1-CAGGS-eGFP was a gift from Rudolf Jaenisch (Addgene plasmid # 22212; http://n2t.net/addgene:22212; RRID:Addgene_22212) [44]. Three mutations were added to the sequence: BglII site 877 killed (now agttct), stopcodon 4278 changed to glycine (now caagggaacggct), and NotI site 5895 killed (now ggggccgc). hAAVS1-1L-TALEN (Addgene plasmid # 35431; http://n2t.net/addgene:35431; RRID:Addgene_35431) and hAAVS1-1R-TALEN plasmids (Addgene plasmid # 35432; http://n2t.net/addgene:35432; RRID:Addgene_35432) were gifts from Feng Zhang [45]. Human *GLIS2* (NM_032575) was cloned from cDNA derived from human podocytes and subsequently inserted into the AAVS1-CAGGS-eGFP, resulting in an N-terminally GFP-tagged fusion protein. Additionally, two mutant cDNAs were inserted into the AAVS1-CAGGS-eGFP plasmid. GLIS2[IVS+1G>T] contains the cDNA of coding exons 1-5 plus the sequence of intron 5 up to the first stop codon resulting in a 341 amino acid long truncated protein containing GLIS2[1–258] and 83 additional residues. For GLIS2[1–121] mutant, the first 121 amino acids of GLIS2 were used. Furthermore, human *GLIS2* was cloned into a modified pcDNA6 vector (Invitrogen) containing an N-terminal GFP-tag. Mouse *Glis2* (NM_031184) was cloned from cDNA from cDNA derived from mouse kidney tissue in a modified pcDNA6 vector (Invitrogen) containing an N-terminal FLAG-tag. Mouse *Glis2* was cloned from cDNA derived from mouse kidney tissue either from wildtype or *Glis2*^Y122X^ animals into a modified pcDNA6 vector containing an N-terminal FLAG-tag and a C-terminal V5-tag using restriction enzymes. Full length and mutation status were assessed by Sanger sequencing.

### Transfection

Cells were grown up to a confluency of 50% for transfection. Plasmids were transfected in HEK293T cells by calcium-chloride transfection. siRNA was transfected in mIMCD3 cells using Lipofectamine RNAimax (Invitrogen) with a 20 nM final concentration according to the manufacturer’s protocol. ON-TARGETplus siRNA SMARTpools (Dharmacon, GE): siControl D-001810-10-20 and siGlis2 (mouse) L-042208-01-0005.

### Polyclonal single-copy transgenic TALEN cell lines

For stable integration into the human AAVS1 locus, HEK293T cells were co-transfected with either of the donor plasmids (AAVS1-CAGGS-eGFP.GLIS2, AAVS1-CAGGS-eGFP.eGFP, AAVS1-CAGGS-eGFP.GLIS2[IVS5+1G>T], AAVS1-CAGGS-eGFP.GLIS2[1–121]) and both hAAVS1-1L-TALEN and hAAVS1-1R-TALEN plasmids by calcium-chloride transfection. On the next day, cells were selected with Puromycin (4 µg/ml, Invitrogen). Successful integration was assessed by genomic PCR. Expression of transgenes was evaluated by live cell imaging.

### Comet assay

mIMCD3 cells were used to perform comet assays after siRNA transfection. 400 nM APH or an equal amount of DMSO was added to the samples 18 h prior to the assay. Single-cell gel electrophoresis was performed using Comet Assay® Kit (Trevigen) according to the manufacturer’s protocol. Images were acquired with an Axiovert 200 M microscope Plan Apochromat 20x / 0.8 DIC objective (all from Carl Zeiss MicroImaging GmbH). Comets were quantified by the OpenComet plugin for Fiji/ImageJ [46]. Before analysis, the original zvi-images were converted to brightness adjusted 8-bit tif-files using Fiji/ImageJ software. The following settings were used in OpenComet: Head finding method: brightest region. Background correction for comet finding was enabled.

### Local laser ablation

Either HEK293T cells, with overexpressed FLAG.GFP or GFP.GLIS2 in Hepes-buffered medium were exposed to local UV-C irradiation on a Zeiss LSM510 confocal microscope (Zen software) equipped with a pulsed 266 nm laser source (DPSL-266, Rapp OptoElectronic GmbH). A special adapter as well as quartz coverslips (SPI supplies, 01019T-AB) and a 100x quartz objective were used to laser ablate the sample. In additional experiments, cells were exposed to local 800 nm laser irradiation on a TCS SP8 MP-OPO (Leica) microscope (Las X software) using a HCX IRAPO L 25x/0.95 W objective. Besides nuclei, cytoplasmic and non-cellular regions were also irradiated as controls for nuclear recruitment.

### Chromatin fractionation

Either GFP.GLIS2 or FLAG.GFP was transiently overexpressed in HEK293T cells. Cells were irradiated with 20 J/m^2^ UV-C. Chromatin fractionation was performed as previously described [12]. Approximately 10^7^ cells were used. The chromatin fraction was sheared by sonication. The fractions were resuspended in Laemmli sample buffer and analyzed by Western blotting. The fractions were analyzed by immunoblotting against GFP (Santa Cruz sc-9996), [1-tubulin (Santa Cruz sc-9104), PCNA (Calbiochem NA03), and H2AX (Cell Signaling 2595).

### Interactome

For each sample one 10-cm dish HEK293T cells transfected with either FLAG.GLIS2 or FLAG.GFP was lysed in IP buffer (20 mM Tris pH7.5, 1% (v/v) TritonX-100, 50 mM NaCl, 25 mM Na_4_P_2_O_7_, 15 mM NaF) supplemented with protease and phosphatase inhibitors (44 µg/ml PMSF, 2 mM Na3VO4, 1x cOmplete™ Protease Inhibitor Cocktail). After sonication, the lysates were first centrifuged 30 min at 14.000 g at 4°C followed by ultracentrifugation for 45 min at 100.000 g and 4°C. The lysates were incubated for 2 h with anti-FLAG (M2) antibody-coated sepharose beads (bimake.com B23102) at 4°C. After three washing steps the beads were incubated with 5% SDS in PBS for 5 min at 95 °C, followed by reduction with 10 mM dithiothreitol and alkylation with 40 mM chloroacetamide. Four biological replicates were performed.

Samples were analyzed by the CECAD Proteomics Facility on an Orbitrap Exploris 480 (Thermo Scientific, granted by the German Research Foundation under INST 1856/71-1 FUGG) mass spectrometer equipped with a FAIMSpro differential ion mobility device that was coupled to an Vanquish neo in trap-and-elute setup (Thermo Scientific). Samples were loaded onto a precolumn (Acclaim 5μm PepMap 300 μ Cartridge) with a flow of 60 μl/min before being reverse-flushed onto an in-house packed analytical column (30 cm length, 75 μm inner diameter, filled with 2.7 μm Poroshell EC120 C18, Agilent). Peptides were chromatographically separated with an initial flow rate of 400 nL/min and the following gradient: initial 2% B (0.1% formic acid in 80 % acetonitrile), up to 6 % in 3 min. Then, flow was reduced to 300 nl/min and B increased to 20% B in 26 min, up to 35% B within 15 min and up to 98% solvent B within 1.0 min while again increasing the flow to 400 nl/min, followed by column wash with 98% solvent B and reequilibration to initial condition. The FAIMS pro was operated at −50V compensation voltage and electrode temperatures of 99.5 °C for the inner and 85 °C for the outer electrode. The mass spectrometer was operated in data-dependent acquisition top 24 mode with MS1 scans acquired from 350 m/z to 1400 m/z at 60k resolution and an AGC target of 300%. MS2 scans were acquired at 15 k resolution with a maximum injection time of 22 ms and an AGC target of 300% in a 1.4 Th window and a fixed first mass of 110 m/z. All MS1 scans were stored as profile, all MS2 scans as centroid.

All mass spectrometric raw data were processed with Maxquant (version 2.4, [47]) using default parameters against the Uniprot HUMAN canonical database (UP5640) merged with the sequence of the FLAG.GLIS2 with the match-between-runs option enabled between replicates. Follow-up analysis was done in Perseus 1.6.15 [47]. Protein groups were filtered for potential contaminants and insecure identifications. Remaining IDs were filtered for data completeness in at least one group and missing values imputed by sigma downshift (0.3 σ width, 1.8 σ downshift). Afterwards, FDR-controlled two-sided t-tests were performed. The resulting identifiers were used for down-stream annotation and further analysis.

### Nano-liquid-chromatography-(nLC)-MS/MS proteomic analysis

After incubation of the eluate with 5 mM DTT and 40 mM CAA, each sample was incubated at RT overnight with trypsin for digestion of the proteins. Digestion was stopped by addition of 0.5% formic acid and further cleaned and desalted using stop-and-go extraction tips (Stagetips) [48]. Prior to MS/MS analysis, peptides were resuspended in 0.1% formic acid and peptides were fractionated by nLC with a 1 hour gradient and a binary buffer system. Samples were finally measured using a quadrupol-orbitrap based QExacutive mass spectrometer (Thermo Scientific) [49].

### Statistics

An unpaired Student’s 2-tailed t-test was used to calculate statistical significance by using GraphPad Prism version 10 (GraphPad Software, San Diego, CA). Statistical significance is represented as follows: *P < 0.05, **P < 0.01, and ***P < 0.001. The results displayed are an average of at least 3 independent experiments or biological replicates, and error bars indicate SEM in all plots unless otherwise indicated.

## Supporting information

Sequences

GLIS2 Interactors

## Data availability

The mass spectrometry proteomics data have been deposited to the ProteomeXchange Consortium via the PRIDE [50] partner repository with the dataset identifier PXD057293. Login Details on request.

## Author contributions

The research was designed by LKE, GGS, and BS. LKE, LS, LEF, MJ, CD, and GGS conducted the experiments and acquired the data. LKE and GGS analyzed the data. MCL provided reagents. LKE, GGS, TB, and BS wrote and edited the manuscript. All authors approved the final version of the paper.

## Acknowledgements

We thank Stefanie Keller, Serena Greco-Torres and Martyna Brütting for their excellent technical assistance. We thank the CECAD Proteomics Facility, especially Jan-Wilm Lackmann, as well as the CECAD Imaging Facility, especially Christian Juengst, and the CECAD *in vivo* Research Facility, especially Simon Tröder, for their technical support.

## Supplemental material

**Fig. S1: Validation and characterization of *Glis2^Y122X^* mice.**

**Fig. S2: Generation and validation of stable GFP.GLIS2 cell lines.**

**Table S1: List of sequences**

**Table S2: List of GLIS2 interactors**

**Movie S1: GFP.GLIS2 recruits to sites of 800 nm laser-induced DNA damage.**

**Movie S2: GFP does not recruit to sites of 800 nm laser-induced DNA damage.**

## Abbreviations

ADPKD: autosomal dominant polycystic kidney disease
DD: DNA damage
DDR: DNA damage response
DSB: DNA strand breaks
GO: gene ontology
mIMCD3: murine inner medulla collecting duct cells
NPH: Nephronophthisis
PAS: periodic acid-Schiff
qPCR: quantitative polymerase chain reaction
ssODN: single-stranded donor oligonucleotide
γH2AX: phosphorylated H2AX

## References

1. Srivastava S, Molinari E, Raman S, Sayer JA. Many Genes—One Disease? Genetics of Nephronophthisis (NPHP) and NPHP-Associated Disorders. Front Pediatr. 2018;5. doi:10.3389/fped.2017.00287

2. Hildebrandt F, Otto E, Rensing C, Nothwang HG, Vollmer M, Adolphs J, et al. A novel gene encoding an SH3 domain protein is mutated in nephronophthisis type 1. Nat Genet. 1997;17: 149–153. doi:10.1038/ng1097-149

3. Hurd TW, Otto EA, Mishima E, Gee HY, Inoue H, Inazu M, et al. Mutation of the Mg2+ transporter SLC41A1 results in a nephronophthisis-like phenotype. J Am Soc Nephrol. 2013;24: 967–977. doi:10.1681/ASN.2012101034

4. Utsch B, Sayer JA, Attanasio M, Pereira RR, Eccles M, Hennies H-C, et al. Identification of the first AHI1 gene mutations in nephronophthisis-associated Joubert syndrome. Pediatr Nephrol. 2006;21: 32–35. doi:10.1007/s00467-005-2054-y

5. Sang L, Miller JJ, Corbit KC, Giles RH, Brauer MJ, Otto EA, et al. Mapping the Nephronophthisis-Joubert-Meckel-Gruber Protein Network Reveals Ciliopathy Disease Genes and Pathways. Cell. 2011;145: 513–528. doi:10.1016/j.cell.2011.04.019

6. Braun DA, Schueler M, Halbritter J, Gee HY, Porath JD, Lawson JA, et al. Whole exome sequencing identifies causative mutations in the majority of consanguineous or familial cases with childhood-onset increased renal echogenicity. Kidney Int. 2016;89: 468–475. doi:10.1038/ki.2015.317

7. Hildebrandt F, Attanasio M, Otto E. Nephronophthisis: Disease Mechanisms of a Ciliopathy. J Am Soc Nephrol. 2009;20: 23–35. doi:10.1681/ASN.2008050456

8. Pazour GJ, Bloodgood RA. Chapter 5 Targeting Proteins to the Ciliary Membrane. Current Topics in Developmental Biology. Academic Press; 2008. pp. 115–149. doi:10.1016/S0070-2153(08)00805-3

9. Chaki M, Airik R, Ghosh AK, Giles RH, Chen R, Slaats GG, et al. Exome capture reveals ZNF423 and CEP164 mutations, linking renal ciliopathies to DNA damage response signaling. Cell. 2012;150: 533–548. doi:10.1016/j.cell.2012.06.028

10. Airik R, Slaats GG, Guo Z, Weiss A-C, Khan N, Ghosh A, et al. Renal-Retinal Ciliopathy Gene Sdccag8 Regulates DNA Damage Response Signaling. J Am Soc Nephrol. 2014;25: 2573–2583. doi:10.1681/ASN.2013050565

11. Slaats GG, Saldivar JC, Bacal J, Zeman MK, Kile AC, Hynes AM, et al. DNA replication stress underlies renal phenotypes in CEP290-associated Joubert syndrome. J Clin Invest. 2015;125: 3657–3666. doi:10.1172/JCI80657

12. Choi HJC, Lin J-R, Vannier J-B, Slaats GG, Kile AC, Paulsen RD, et al. NEK8 links the ATR-regulated replication stress response and S phase CDK activity to renal ciliopathies. Mol Cell. 2013;51: 423–439. doi:10.1016/j.molcel.2013.08.006

13. Slaats GG, Giles RH. Are renal ciliopathies (replication) stressed out? Trends Cell Biol. 2015;25: 317–319. doi:10.1016/j.tcb.2015.03.005

14. Vasanth S, ZeRuth G, Kang HS, Jetten AM. Identification of nuclear localization, DNA binding, and transactivating mechanisms of Kruppel-like zinc finger protein Gli-similar 2 (Glis2). J Biol Chem. 2011;286: 4749–4759. doi:10.1074/jbc.M110.165951

15. Attanasio M, Uhlenhaut NH, Sousa VH, O’Toole JF, Otto E, Anlag K, et al. Loss of GLIS2 causes nephronophthisis in humans and mice by increased apoptosis and fibrosis. Nat Genet. 2007;39: 1018–1024. doi:10.1038/ng2072

16. Kim Y-S, Kang HS, Herbert R, Beak JY, Collins JB, Grissom SF, et al. Krüppel-Like Zinc Finger Protein Glis2 Is Essential for the Maintenance of Normal Renal Functions. Mol Cell Biol. 2008;28: 2358–2367. doi:10.1128/MCB.01722-07

17. Lu D, Rauhauser A, Li B, Ren C, McEnery K, Zhu J, et al. Loss of Glis2/NPHP7 causes kidney epithelial cell senescence and suppresses cyst growth in the Kif3a mouse model of cystic kidney disease. Kidney Int. 2016;89: 1307–1323. doi:10.1016/j.kint.2016.03.006

18. Zhang F, Nakanishi G, Kurebayashi S, Yoshino K, Perantoni A, Kim Y-S, et al. Characterization of Glis2, a Novel Gene Encoding a Gli-related, Krüppel-like Transcription Factor with Transactivation and Repressor Functions Roles in Kidney Development and Neurogenesis. J Biol Chem. 2002;277: 10139–10149. doi:10.1074/jbc.M108062200

19. Halbritter J, Porath JD, Diaz KA, Braun DA, Kohl S, Chaki M, et al. Identification of 99 novel mutations in a worldwide cohort of 1,056 patients with a nephronophthisis-related ciliopathy. Hum Genet. 2013;132: 865–884. doi:10.1007/s00439-013-1297-0

20. Al Alawi I, Powell L, Rice SJ, Al Riyami MS, Al-Riyami M, Al Salmi I, et al. Case Report: A Novel In-Frame Deletion of GLIS2 Leading to Nephronophthisis and Early Onset Kidney Failure. Front Genet. 2021;12: 791495. doi:10.3389/fgene.2021.791495

21. Snyder RD, Regan JD. Aphidicolin inhibits repair of DNA in UV-irradiated human fibroblasts. Biochemical and Biophysical Research Communications. 1981;99: 1088–1094. doi:10.1016/0006-291X(81)90730-0

22. Lee YY, Yu YB, Gunawardena HP, Xie L, Chen X. BCLAF1 is a radiation-induced H2AX-interacting partner involved in γH2AX-mediated regulation of apoptosis and DNA repair. Cell Death Dis. 2012;3: e359–e359. doi:10.1038/cddis.2012.76

23. Ray Chaudhuri A, Nussenzweig A. The multifaceted roles of PARP1 in DNA repair and chromatin remodelling. Nat Rev Mol Cell Biol. 2017;18: 610–621. doi:10.1038/nrm.2017.53

24. Szklarczyk D, Kirsch R, Koutrouli M, Nastou K, Mehryary F, Hachilif R, et al. The STRING database in 2023: protein-protein association networks and functional enrichment analyses for any sequenced genome of interest. Nucleic Acids Res. 2023;51: D638–D646. doi:10.1093/nar/gkac1000

25. Ahel D, Hořejší Z, Wiechens N, Polo SE, Garcia-Wilson E, Ahel I, et al. Poly(ADP-ribose)– Dependent Regulation of DNA Repair by the Chromatin Remodeling Enzyme ALC1. Science. 2009;325: 1240–1243. doi:10.1126/science.1177321

26. Smith R, Sellou H, Chapuis C, Huet S, Timinszky G. CHD3 and CHD4 recruitment and chromatin remodeling activity at DNA breaks is promoted by early poly(ADP-ribose)-dependent chromatin relaxation. Nucleic Acids Res. 2018;46: 6087–6098. doi:10.1093/nar/gky334

27. Cargill MJ, Morales A, Ravishankar S, Warren EH. RNA helicase, DDX3X, is actively recruited to sites of DNA damage in live cells. DNA Repair (Amst). 2021;103: 103137. doi:10.1016/j.dnarep.2021.103137

28. Furuta T, Takemura H, Liao Z-Y, Aune GJ, Redon C, Sedelnikova OA, et al. Phosphorylation of Histone H2AX and Activation of Mre11, Rad50, and Nbs1 in Response to Replication-dependent DNA Double-strand Breaks Induced by Mammalian DNA Topoisomerase I Cleavage Complexes *. Journal of Biological Chemistry. 2003;278: 20303–20312. doi:10.1074/jbc.M300198200

29. Essers J, Theil AF, Baldeyron C, van Cappellen WA, Houtsmuller AB, Kanaar R, et al. Nuclear Dynamics of PCNA in DNA Replication and Repair. Mol Cell Biol. 2005;25: 9350–9359. doi:10.1128/MCB.25.21.9350-9359.2005

30. Dinant C, de Jager M, Essers J, van Cappellen WA, Kanaar R, Houtsmuller AB, et al. Activation of multiple DNA repair pathways by sub-nuclear damage induction methods. J Cell Sci. 2007;120: 2731–2740. doi:10.1242/jcs.004523

31. Polo SE, Kaidi A, Baskcomb L, Galanty Y, Jackson SP. Regulation of DNA-damage responses and cell-cycle progression by the chromatin remodelling factor CHD4. The EMBO Journal. 2010;29: 3130–3139. doi:10.1038/emboj.2010.188

32. Yao J, Lei P-J, Li Q-L, Chen J, Tang S-B, Xiao Q, et al. GLIS2 promotes colorectal cancer through repressing enhancer activation. Oncogenesis. 2020;9: 1–15. doi:10.1038/s41389-020-0240-1

33. Jetten AM. GLIS1-3 transcription factors: critical roles in the regulation of multiple physiological processes and diseases. Cell Mol Life Sci. 2018;75: 3473–3494. doi:10.1007/s00018-018-2841-9

34. Zhang C, Rehman M, Tian X, Pei SLC, Gu J, Bell TA, et al. Glis2 is an early effector of polycystin signaling and a target for therapy in polycystic kidney disease. Nat Commun. 2024;15: 3698. doi:10.1038/s41467-024-48025-6

35. Pogue RE, Cavalcanti DP, Shanker S, Andrade RV, Aguiar LR, de Carvalho JL, et al. Rare genetic diseases: update on diagnosis, treatment and online resources. Drug Discov Today. 2018;23: 187–195. doi:10.1016/j.drudis.2017.11.002

36. Stokman MF, Saunier S, Benmerah A. Renal Ciliopathies: Sorting Out Therapeutic Approaches for Nephronophthisis. Front Cell Dev Biol. 2021;9: 653138. doi:10.3389/fcell.2021.653138

37. Garcia H, Serafin AS, Silbermann F, Porée E, Viau A, Mahaut C, et al. Agonists of prostaglandin E2 receptors as potential first in class treatment for nephronophthisis and related ciliopathies. Proceedings of the National Academy of Sciences. 2022;119: e2115960119. doi:10.1073/pnas.2115960119

38. Truett GE, Heeger P, Mynatt RL, Truett AA, Walker JA, Warman ML. Preparation of PCR-quality mouse genomic DNA with hot sodium hydroxide and tris (HotSHOT). Biotechniques. 2000;29: 52, 54. doi:10.2144/00291bm09

39. Ran FA, Hsu PD, Wright J, Agarwala V, Scott DA, Zhang F. Genome engineering using the CRISPR-Cas9 system. Nat Protoc. 2013;8: 2281–2308. doi:10.1038/nprot.2013.143

40. Wang H, Yang H, Shivalila CS, Dawlaty MM, Cheng AW, Zhang F, et al. One-step generation of mice carrying mutations in multiple genes by CRISPR/Cas-mediated genome engineering. Cell. 2013;153: 910–918. doi:10.1016/j.cell.2013.04.025

41. Wefers B, Bashir S, Rossius J, Wurst W, Kühn R. Gene editing in mouse zygotes using the CRISPR/Cas9 system. Methods. 2017;121–122: 55–67. doi:10.1016/j.ymeth.2017.02.008

42. Kieckhöfer E, Slaats GG, Ebert LK, Albert M-C, Dafinger C, Kashkar H, et al. Primary cilia suppress Ripk3-mediated necroptosis. Cell Death Discov. 2022;8: 1–12. doi:10.1038/s41420-022-01272-2

43. Jain M, Kaiser RWJ, Bohl K, Hoehne M, Göbel H, Bartram MP, et al. Inactivation of Apoptosis Antagonizing Transcription Factor in tubular epithelial cells induces accumulation of DNA damage and nephronophthisis. Kidney Int. 2019;95: 846–858. doi:10.1016/j.kint.2018.10.034

44. Hockemeyer D, Soldner F, Beard C, Gao Q, Mitalipova M, DeKelver RC, et al. Efficient targeting of expressed and silent genes in human ESCs and iPSCs using zinc-finger nucleases. Nat Biotechnol. 2009;27: 851–857. doi:10.1038/nbt.1562

45. Sanjana NE, Cong L, Zhou Y, Cunniff MM, Feng G, Zhang F. A transcription activator-like effector toolbox for genome engineering. Nat Protoc. 2012;7: 171–192. doi:10.1038/nprot.2011.431

46. Gyori BM, Venkatachalam G, Thiagarajan PS, Hsu D, Clement M-V. OpenComet: an automated tool for comet assay image analysis. Redox Biol. 2014;2: 457–465. doi:10.1016/j.redox.2013.12.020

47. Tyanova S, Temu T, Sinitcyn P, Carlson A, Hein MY, Geiger T, et al. The Perseus computational platform for comprehensive analysis of (prote)omics data. Nat Methods. 2016;13: 731–740. doi:10.1038/nmeth.3901

48. Rappsilber J, Ishihama Y, Mann M. Stop and go extraction tips for matrix-assisted laser desorption/ionization, nanoelectrospray, and LC/MS sample pretreatment in proteomics. Anal Chem. 2003;75: 663–670. doi:10.1021/ac026117i

49. Michalski A, Damoc E, Hauschild J-P, Lange O, Wieghaus A, Makarov A, et al. Mass spectrometry-based proteomics using Q Exactive, a high-performance benchtop quadrupole Orbitrap mass spectrometer. Mol Cell Proteomics. 2011;10: M111.011015. doi:10.1074/mcp.M111.011015

50. Perez-Riverol Y, Bai J, Bandla C, García-Seisdedos D, Hewapathirana S, Kamatchinathan S, et al. The PRIDE database resources in 2022: a hub for mass spectrometry-based proteomics evidences. Nucleic Acids Res. 2022;50: D543–D552. doi:10.1093/nar/gkab1038

