## Supplementary material for "The nephronophthisis protein *GLIS2*/*NPHP7* is required for the DNA damage response in kidney tubular epithelial cells": Suppl. Fig. 1 & 2

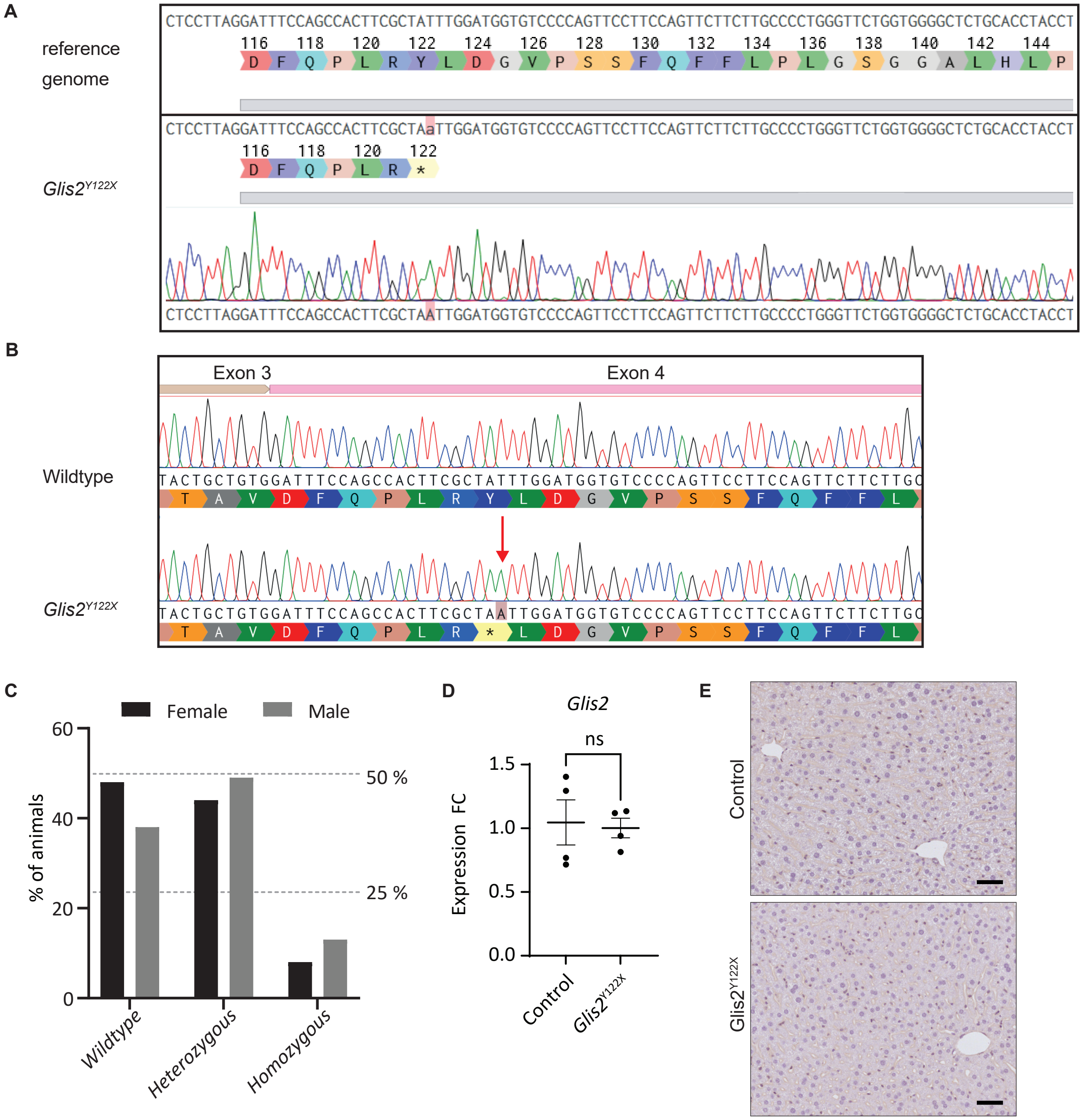


**Figure S1. Validation and characterization of *Glis2*$^{Y122X}$ mice.** (a) Sanger sequencing of gDNA isolated from tail tissue from a homozygous *Glis2*$^{Y122X}$ mouse confirms introduction of the point mutation. (b) Sanger sequencing of cDNA isolated from kidney tissue from a homozygous *Glis2*$^{Y122X}$ mouse confirms the introduction of the point mutation. (c) Genotype distribution in litters born from experimental matings (*Glis2*$^{WT/Y122X}$ x *Glis2*$^{WT/Y122X}$). 94 mice were analyzed. Expected mendelian ratios are indicated with dotted lines (50 % for heterozygous (*Glis2*$^{WT/Y122X}$) and 25 % for wildtype (*Glis2*$^{WT/WT}$) and homozygous (*Glis2*$^{Y122X/Y122X}$)). *Glis2*$^{Y122X}$ mice show a non-Mendelian pattern of inheritance with fewer homozygous mutant mice born than the 25% expected when crossing two heterozygous animals. (d) Quantitative polymerase chain reaction (qPCR) to assess *Glis2* mRNA expression in *Glis2*$^{Y122X}$ mice (N=4). No significant difference in mRNA expression of mutant *Glis2* was revealed, indicating that the point mutation did not induce nonsense-mediated RNA decay. (e) yH2A.X staining of liver tissue of 16-week-old *Glis2*$^{Y122X}$ and control mice (scale bars: 50 µm, N=4).

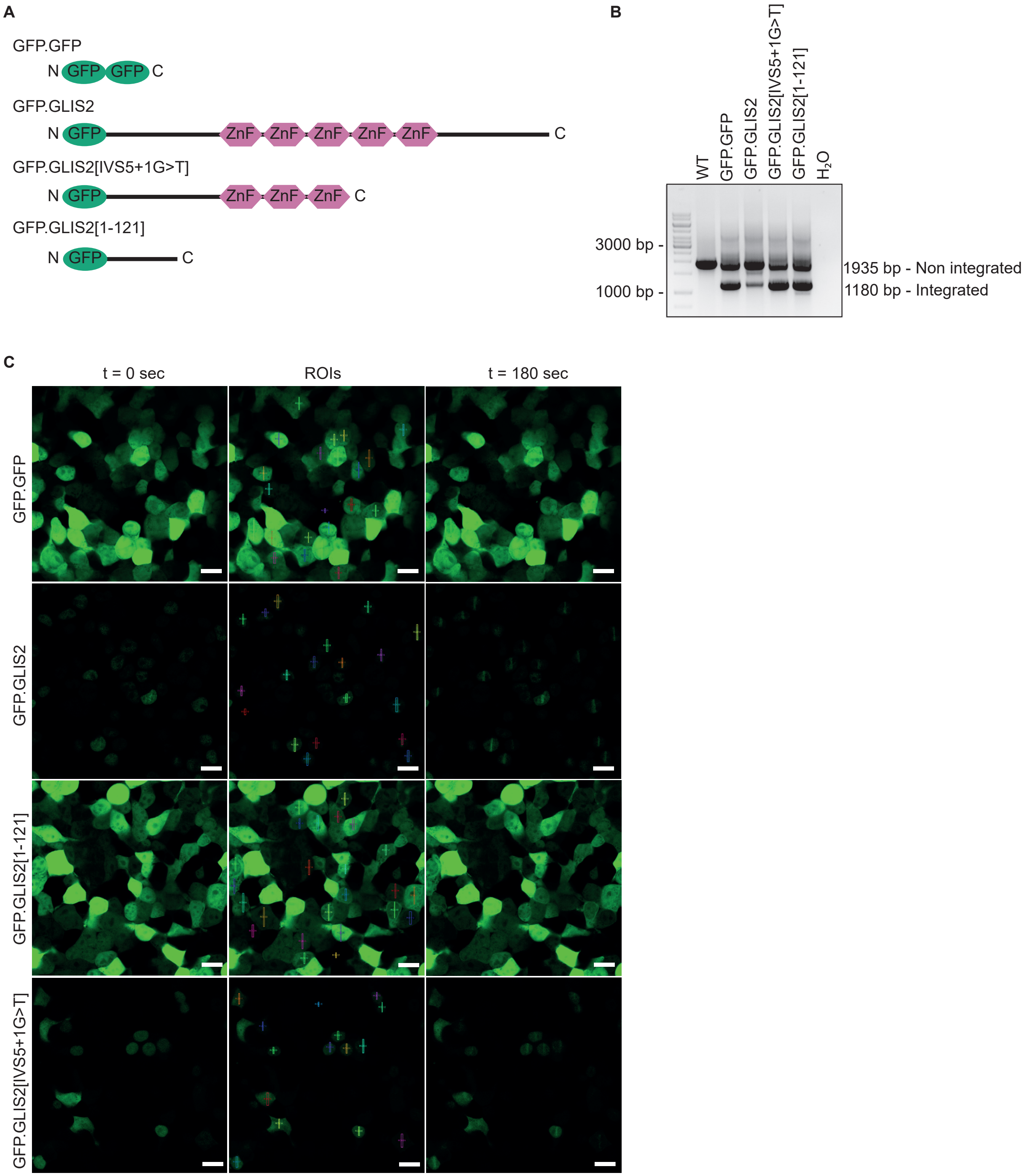


**Fig S2. Generation and validation of stable GFP.GLIS2 cell lines.** (a) Schematic overview of the stable GFP-tagged wildtype and mutant GLIS2 TALEN HEK293T cell lines. (b) Integration PCR showing the integration of the GFP.GFP and GFP.GLIS2 wildtype and mutant constructs in the AAVS1 locus. (c) GFP-tagged GLIS2 variant kinetics after 800 nm laser irradiation. Redistribution of protein was follow over time (end t = 180 s, scale bar = 10 μm, N=3)
